# *Pseudomonas aeruginosa* isolates co-incubated with *Acanthamoeba castellanii* exhibit phenotypes similar to chronic cystic fibrosis isolates

**DOI:** 10.1101/2020.02.25.964320

**Authors:** Wai Leong, Carla Lutz, Jonathan Williams, Yan Hong Poh, Benny Yeo Ken Yee, Cliff Chua, Scott A. Rice, Michael Givskov, Martina Sanderson-Smith, Diane McDougald

## Abstract

The opportunistic pathogen, *Pseudomonas aeruginosa*, is ubiquitous in the environment, and in humans is capable of causing acute and chronic infections. *P. aeruginosa*, when co-incubated with the bacterivorous amoeba, *Acanthamoeba castellanii*, for extended periods, produced genetic and phenotypic variants. Sequencing of late-stage amoeba-adapted *P. aeruginosa* isolates demonstrated single nucleotide polymorphisms within genes that encode known virulence factors, and this correlated with a reduction in expression of virulence traits. Virulence towards the nematode, *Caenorhabditis elegans*, was attenuated in late-stage amoeba-adapted *P. aeruginosa* compared to early stage amoeba-adapted and non-adapted counterparts. Late-stage amoeba-adapted *P. aeruginosa* lost competitive fitness compared to non-adapted counterparts when grown in nutrient rich media. However, non-adapted *P. aeruginosa* were rapidly cleared by amoeba predation, whereas late-stage amoeba-adapted isolates remained in higher numbers 24 h after ingestion by amoeba. In addition, there was reduced uptake by macrophage of amoeba-adapted isolates and reduced uptake by human neutrophils as well as increased survival in the presence of neutrophils. Our findings indicate that the selection imposed by amoeba on *P. aeruginosa* resulted in reduced virulence over time. Importantly, the genetic and phenotypic traits possessed by late-stage amoeba-adapted *P. aeruginosa* are similar to what is observed for isolates obtained from chronic cystic fibrosis infections. This notable overlap in adaptation to different host types suggests similar selection pressures among host cell types.

**Author Summary:** *Pseudomonas aeruginosa* is an opportunistic pathogen that causes both acute infections in plants and animals, including humans and also causes chronic infections in immune compromised and cystic fibrosis patients. This bacterium is commonly found in soils and water where bacteria are constantly under threat of being consumed by the bacterial predators, protozoa. To escape being killed, bacteria have evolved a suite of mechanisms that protect them from being consumed or digested. Here we examined the effect of long-term predation on the genotype and phenotypes expressed by *P. aeruginosa.* We show that long-term co-incubation with protozoa resulted in mutations in the bacteria that made them less pathogenic. This is particularly interesting as we see similar mutations arise in bacteria associated with chronic infections. Thus, predation by protozoa and long term colonization of the human host may represent similar environments that select for similar losses in gene functions.

## Introduction

Many virulence traits of microorganisms are regulated in response to the environment in order to invade a host, obtain resources, defend against predation by heterotrophic protists, or establish a replication niche. The evolution of virulence, i.e. harm caused by a pathogen towards its host, is a long-standing subject of investigation with important implications for human health. Most opportunistic pathogens are not transmitted person to person but rather transit through the environment between hosts and therefore, it is unlikely that virulence traits evolve in the host (1–3). Rather, it is more likely that these traits evolve in the environment.

Predation by protists, or protozoa, is a major mortality factor for bacteria in the environment (4). Virulence traits, particularly those that cause human disease, are hypothesized to have evolved in response to and are maintained by predation pressure, which supports the “coincidental evolution” hypothesis. This hypothesis states that virulence is a coincidental consequence of adaptation to other ecological niches (5–7). Coincidental evolution is supported by examples of factors that play roles in both grazing resistance and virulence towards mammalian hosts (7–9), including traits such as cell-surface alterations, increased swimming speed, toxin release and biofilm formation (5, 7). Conversely, virulence traits may be attenuated or lost when organisms adapt to form a more commensal relationship with a host (10–12). Microorganisms may also develop specific virulence traits against a specific host becoming a specialist pathogen. Although there are many hypotheses for how virulence traits evolve, there have been few studies on the adaptation of specific virulence traits to different host types and environments. Such studies are particularly important for understanding how opportunistic pathogens evolve (13). *Pseudomonas aeruginosa* is a versatile opportunistic pathogen found in a wide variety of natural habitats. *P. aeruginosa* has a large (6.3 Mb) genome containing many genes for metabolism and antibiotic resistance (14), and coupled with a complex regulatory network allows it to effectively survive in a variety of niches. *P. aeruginosa* is an important pathogen, responsible for both acute nosocomial infections (15) and chronic infections in leg ulcers and particularly in the lungs of cystic fibrosis (CF) patients (16). In the CF lung, it has been shown to evolve towards a more commensal lifestyle by altering the expression of acute virulence traits such as motility, quorum sensing and toxin production (17). While there are many studies addressing the evolution of *P. aeruginosa* in the CF lung (17–19), there is less known about the impact of protozoan predation on the evolution of virulence. To address this lack of knowledge, this study investigated the adaptation of *P. aeruginosa* during long-term co-incubation with the amoeba predator, *Acanthamoeba castellanii. P. aeruginosa* was co-incubated with *A. castellanii* for 42 days and the impact of co-incubation assessed using a range of phenotypes, including virulence in a *Caenorhabditis elegans* infection model. Adapted populations were also sequenced to investigate the range of mutations that occurred during co-incubation.

## Results

### Genotypic changes in adapted strains

The number of synonymous and non-synonymous single nucleotide polymorphisms (sSNPs and nsSNPs, respectively) occurring in amoeba-adapted and non-adapted populations were determined. There were 54 nsSNPs and 17 sSNPs detected in the 42 d adapted populations and 65 nsSNPs and 19 sSNPs detected in the 42 d non-adapted populations compared to the parent strain. The genes that contained nsSNPs were grouped into gene functions based on Clusters of Orthologous Groups (COGs) and were distributed into 14 COG categories (Fig 1). Mutations in genes involved in transcription occurred in equal numbers in adapted and non-adapted populations. In contrast, nsSNPs in coenzyme metabolism, lipid metabolism, cell motility and secondary metabolite production occurred solely within the amoeba-adapted population, while COGs representing inorganic ion transport and energy production were over-represented in the non-adapted population (Fig 1).

**Fig 1.**
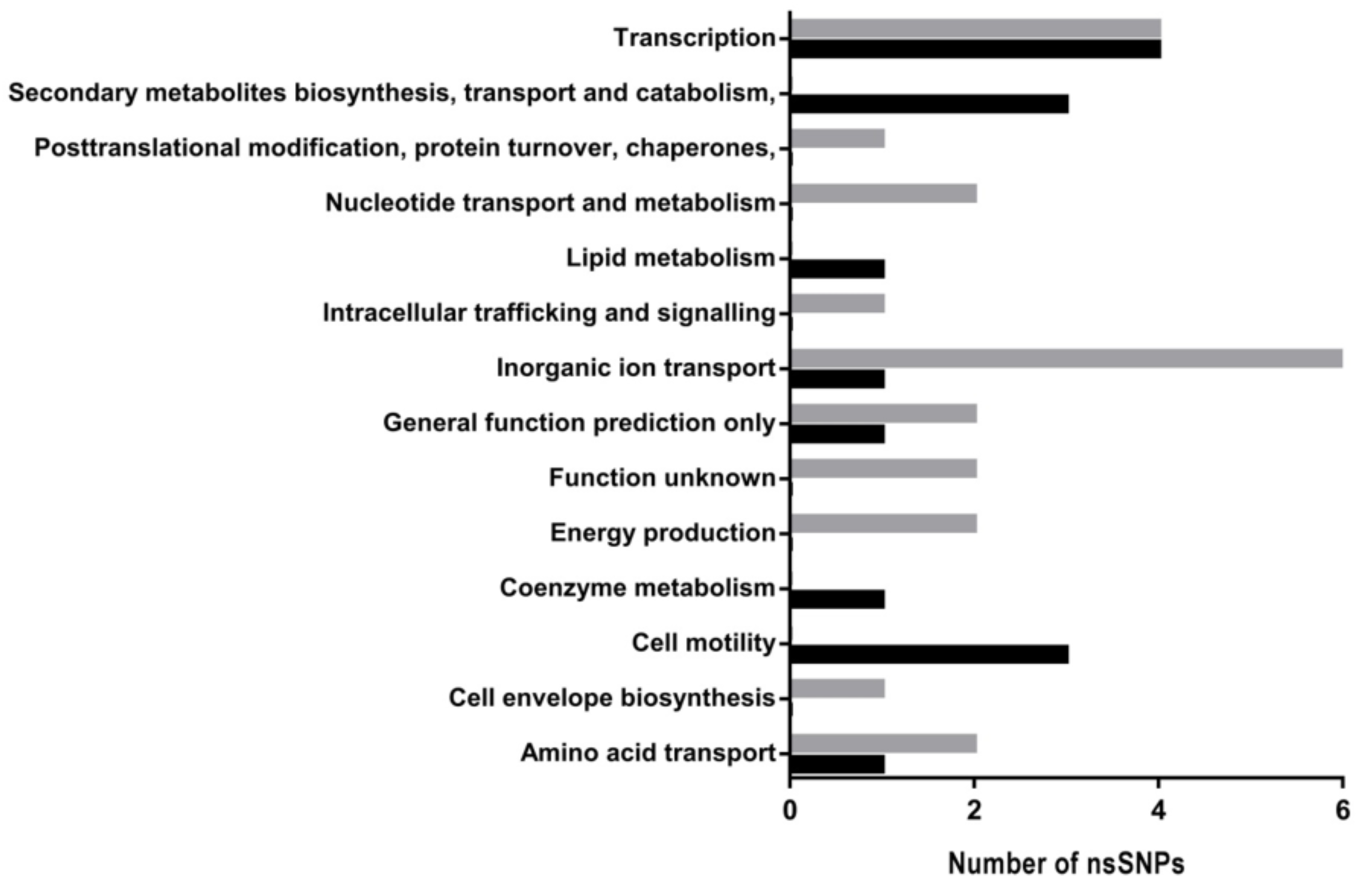
Gene classifications. Classification of genes containing nsSNPs grouped by COG functional class for amoeba-adapted (black bars) and non-adapted (grey bars) populations.

The following nine genes were independently mutated in all three amoeba-adapted replicate populations: the virulence regulator *vreA*, chemotaxis genes *pctB* and PA3349, lipopolysaccharide biosynthetic gene *lpxO2*, *ppiA* and *polA* involved in translation, a cytochrome oxidase *ccpR*, the siderophore *fvbA*, and the hypothetical protein PA3638. The mutations with the highest frequency in the amoeba-adapted populations occurred within motility genes, however mutations in different genes were responsible for the loss of motility observed in the replicate experimental populations. For example, in population 1, 55.17 % of the reads in the *flgF* gene and 34.63 % of the reads in *flgH* contained a nsSNP. Amoeba-adapted populations 2 and 3 contained considerable variation in the gene encoding *flgK*, where 42.31 % of the *flgK* reads from population 2 contained nsSNPs and 96.43 % of *flgK* reads from population 3 contained a gene deletion. In addition to flagellar-mediated motility, all 42 d amoeba-adapted populations contained deletions or SNPs within genes associated with twitching motility, specifically 39.53 % of reads encoding *pilN*, 46.43 % reads encoding *pilM* and 25.53 % of reads encoding *pilT* were detected in population 1, 2 and 3 respectively. Further details about the gene function, frequency of SNPs and codon/amino acid substitution can be found in Table 1.

**Table 1.**
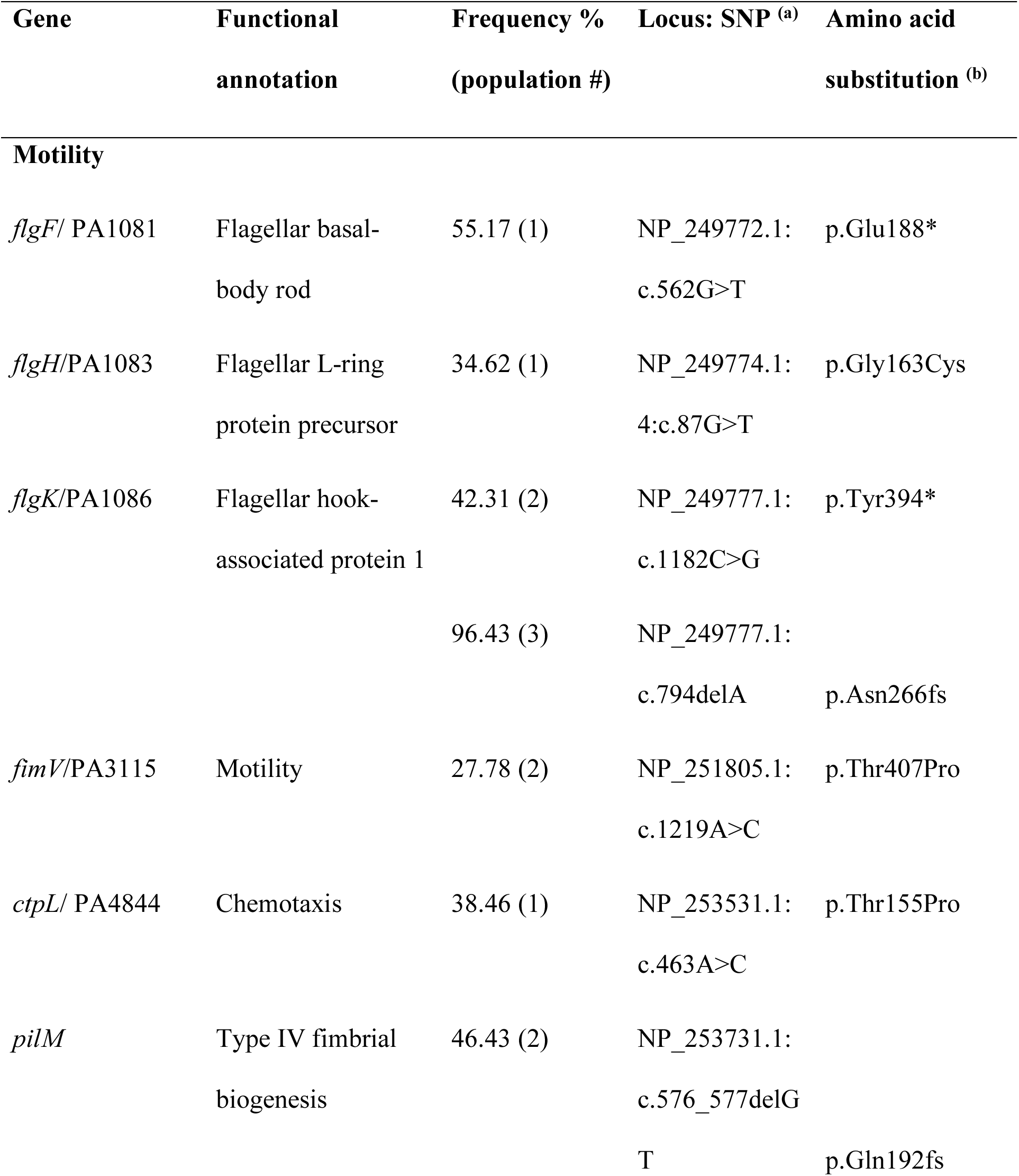
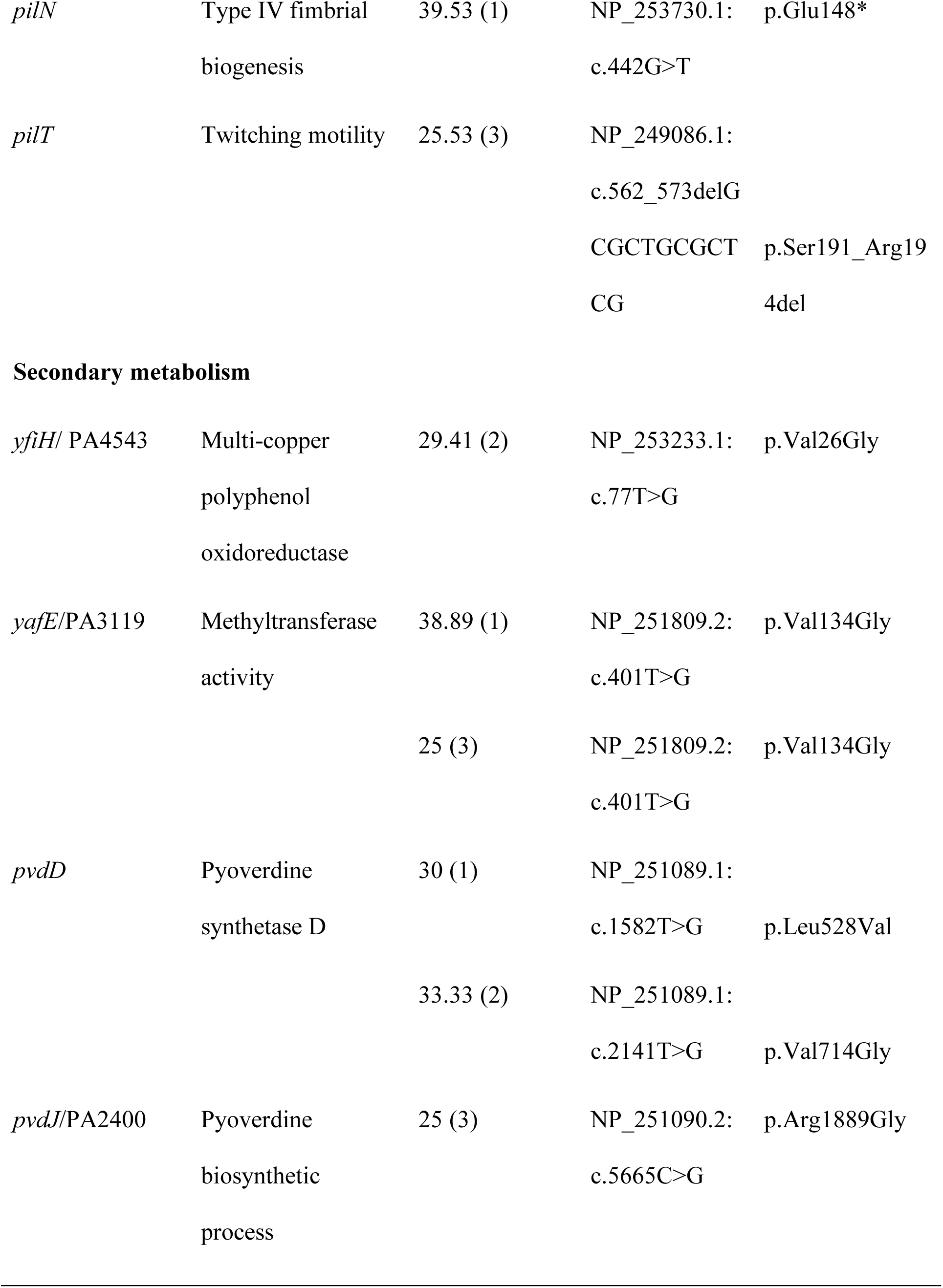
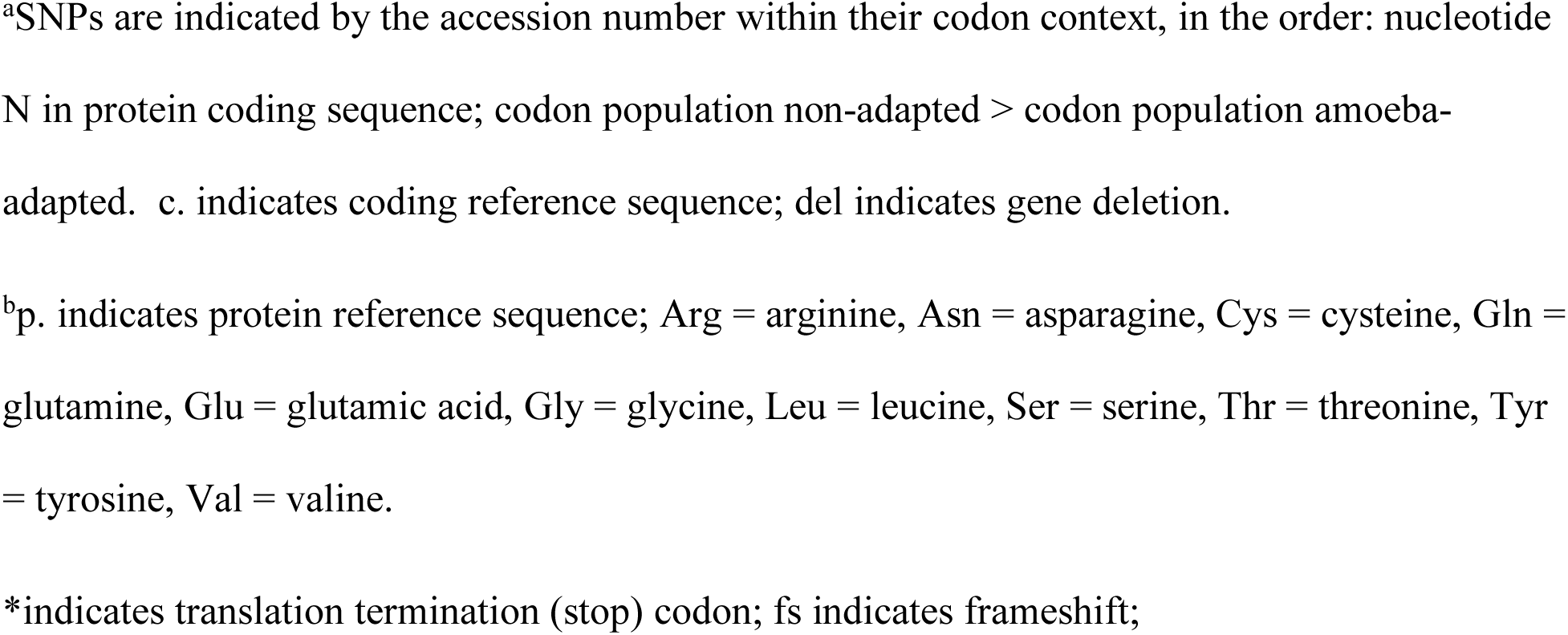
Non-synonymous SNPs and gene deletions in genes associated with motility and secondary metabolism that occurred solely in amoeba-adapted 42 d populations

| Gene | Functional<br>annotation | Frequency %<br>(population #) | Locus: SNP <sup>(a)</sup> | Amino acid<br>substitution <sup>(b)</sup> |
| --- | --- | --- | --- | --- |
| **Motility** |  |  |  |  |
| *flgF*/ PA1081 | Flagellar basal-<br>body rod | 55.17 (1) | NP_249772.1:<br>c.562G>T | p.Glu188* |
| *flgH*/PA1083 | Flagellar L-ring<br>protein precursor | 34.62 (1) | NP_249774.1:<br>4:c.87G>T | p.Gly163Cys |
| *flgK*/PA1086 | Flagellar hook-<br>associated protein 1 | 42.31 (2) | NP_249777.1:<br>c.1182C>G | p.Tyr394* |
|  |  | 96.43 (3) | NP_249777.1:<br>c.794delA | p.Asn266fs |
| *fimV*/PA3115 | Motility | 27.78 (2) | NP_251805.1:<br>c.1219A>C | p.Thr407Pro |
| *ctpL*/ PA4844 | Chemotaxis | 38.46 (1) | NP_253531.1:<br>c.463A>C | p.Thr155Pro |
| *pilM* | Type IV fimbrial<br>biogenesis | 46.43 (2) | NP_253731.1:<br>c.576_577delG<br>T | p.Gln192fs |
| *pilN* | Type IV fimbrial<br>biogenesis | 39.53 (1) | NP_253730.1:<br>c.442G>T | p.Glu148* |
| *pilT* | Twitching motility | 25.53 (3) | NP_249086.1:<br>c.562_573delG<br>CGCTGCGCT<br>CG | p.Ser191_Arg19<br>4del |
| **Secondary metabolism** |  |  |  |  |
| *yfiH/ PA4543* | Multi-copper<br>polyphenol<br>oxidoreductase | 29.41 (2) | NP_253233.1:<br>c.77T>G | p.Val26Gly |
| *yafE/PA3119* | Methyltransferase<br>activity | 38.89 (1) | NP_251809.2:<br>c.401T>G | p.Val134Gly |
|  |  | 25 (3) | NP_251809.2:<br>c.401T>G | p.Val134Gly |
| *pvdD* | Pyoverdine<br>synthetase D | 30 (1) | NP_251089.1:<br>c.1582T>G | p.Leu528Val |
|  |  | 33.33 (2) | NP_251089.1:<br>c.2141T>G | p.Val714Gly |
| *pvdJ/PA2400* | Pyoverdine<br>biosynthetic<br>process | 25 (3) | NP_251090.2:<br>c.5665C>G | p.Arg1889Gly |

<sup>a</sup>SNPs are indicated by the accession number within their codon context, in the order: nucleotide N in protein coding sequence; codon population non-adapted > codon population amoeba-adapted. c. indicates coding reference sequence; del indicates gene deletion. <sup>b</sup>p. indicates protein reference sequence; Arg = arginine, Asn = asparagine, Cys = cysteine, Gln = glutamine, Glu = glutamic acid, Gly = glycine, Leu = leucine, Ser = serine, Thr = threonine, Tyr = tyrosine, Val = valine. \*indicates translation termination (stop) codon; fs indicates frameshift;

Analysis using DAVID functional tools predicted that the second unique biological process affected by SNPs as a result of long-term adaption with amoeba is the biosynthesis, transport and catabolism of secondary metabolites. All 42 d amoeba-adapted populations contain SNPs in *pvdJ* or *pvdD* and these mutations are predicted to affect the synthesis of pyoverdine. Additionally, in an analysis of the 42 d amoeba-adapted populations, *yfiH* and *yafE* were also categorized under secondary metabolite synthesis (20). Non-synonymous SNPs within these genes may further prevent synthesis of secondary metabolites.

### Effect of co-incubation of *P. aeruginosa* with *A. castellanii* on motility

The mutations with the highest frequency in the amoeba-adapted populations occurred within motility genes. Thus, swimming, swarming and twitching motility of the isolates from adapted and non-adapted populations were compared. The long-term co-incubation of *P. aeruginosa* with amoeba resulted in a reduction in twitching motility (Fig 2a; F_2, 534_ = 295.1, *p* < 0.001). *P. aeruginosa* isolates from the 3 d amoeba-adapted and non-adapted populations did not differ significantly (*p* = 0.53), however, after 24 d twitching was significantly reduced compared to the non-adapted isolates (*p* < 0.001) and in the 42 d population, the mean twitching motility of amoeba-adapted isolates was 10-fold less than isolates that were incubated in the absence of *A. castellanii* (*p* < 0.001).

**Fig 2.**
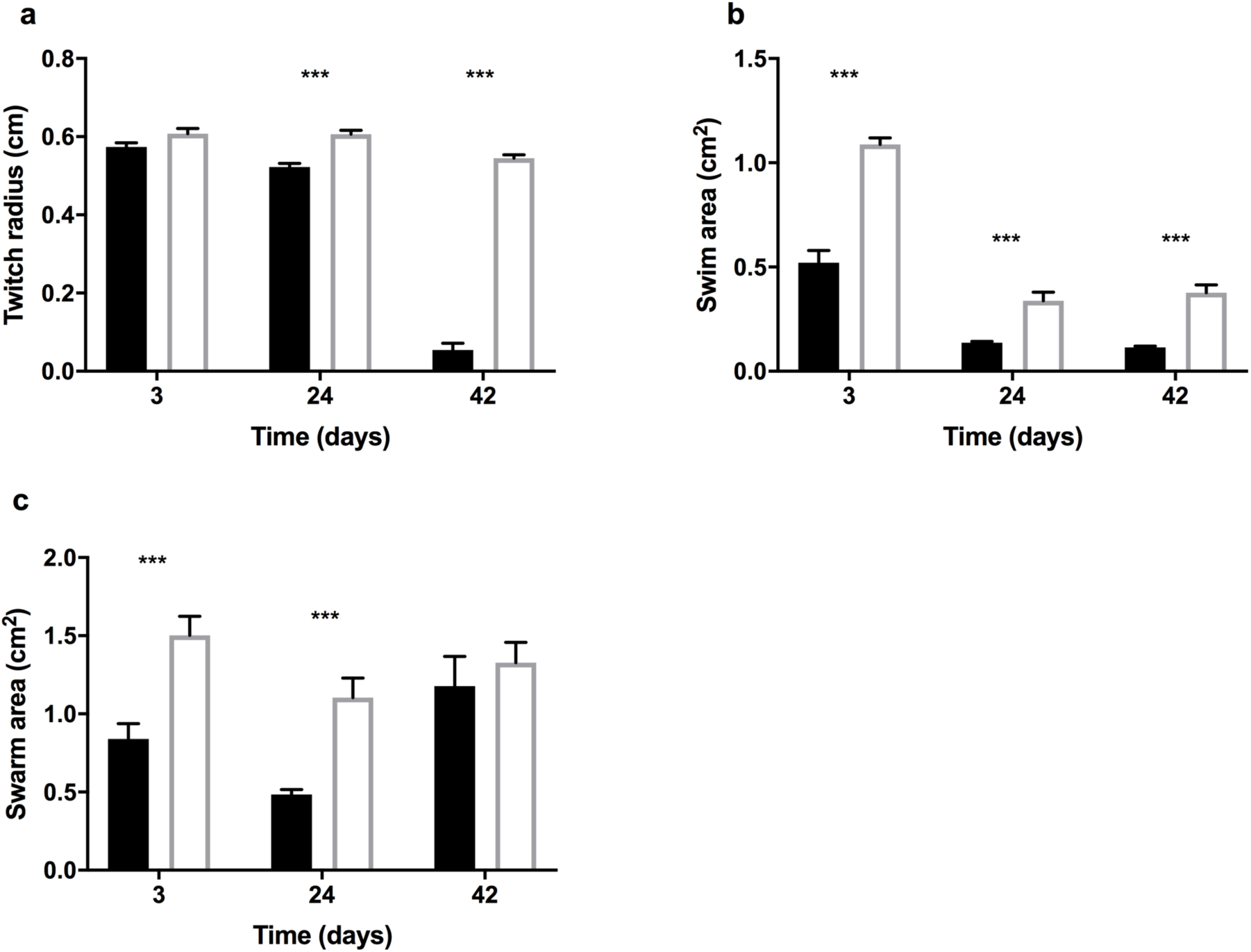
Effect of co-incubation of *P. aeruginosa* with *A. castellanii* on motility. Twitching (a), swimming (b) and swarming motility (c) of *P. aeruginosa* isolates derived from amoeba-adapted (closed) and non-adapted (open) isolates on days 3, 24 and 42. Data are presented as Means ± SEM. *** p < 0.001.

Co-incubation with *A. castellanii* also resulted in a decrease in swimming motility (Fig 2b; *F*_2, 534_ = 15.6, *p* < 0.001), where *P. aeruginosa* isolates from 3 d amoeba-adapted populations had a swim area of half that of isolates from non-adapted populations (*p* < 0.001). This pattern of reduced swimming motility was observed for amoeba-adapted isolates from the 24 and 42 d populations as well (*p* < 0.001 and *p* < 0.001, respectively).

*P. aeruginosa* isolates demonstrated a reduction in swarming motility as a result of co-incubation with amoeba, which varied over time in a non-linear fashion (Fig 2c; *F*_2, 534_ = 7.597, *p* < 0.001). Swarming was significantly reduced in isolates from amoeba-adapted populations at days 3 and 24 (*F*_1, 534_ = 21.73, *p* < 0.001). Post hoc analysis shows that after 3 d of co-incubation the swarming distance of non-adapted isolates of *P. aeruginosa* was twice that of amoeba-adapted isolates (*p* > 0.001). After 24 d of co-incubation the swarming distance exhibited by *P. aeruginosa* isolates derived from amoeba-adapted and non-adapted populations were further reduced, however, there is still a significant reduction in swarming of amoeba-adapted isolates compared to non-adapted isolates (*p<* 0.001). After 42 d there was no significant difference between the average swarming motility of isolates from either population (*p* = 0.189).

### Biofilm formation and planktonic growth of isolates from adapted populations

As flagella and pili also impact biofilm formation, biofilm biomass and planktonic growth rates of adapted and non-adapted isolates were compared. Co-incubation of *P. aeruginosa* with *A. castellanii* had a significant effect on *P. aeruginosa* surface colonization (Fig 3a; *F*_2, 354_ = 15.7, *p* < 0.001). Post hoc analysis revealed no difference between treatments after 3 d (*p =* 0.998). However, *P. aeruginosa* from amoeba-adapted day 24 populations formed 10-fold less biofilm than isolates from non-adapted day 24 populations (*p <* 0.001). Although the average biomass of biofilms formed by the amoeba-adapted isolates increased after 42 d of co-incubation with amoeba, biofilm biomass remained 2-fold lower than that of the non-adapted population (*p* < 0.001). Additionally, the presence of amoeba exerted a strong negative effect on the planktonic growth of *P. aeruginosa* isolates derived from the amoeba-adapted population (Fig 3b; *F*_1, 354_ = 29.6, *p* < 0.001). The planktonic growth rate of *P. aeruginosa* after 3 d was the same regardless of the population (*p* = 0.56). However, after 24 and 42 d of amoebal-driven selection the planktonic growth rate of amoeba-adapted derived isolates was significantly less than the non-adapted isolates (*p* < 0.05 and *p* < 0.001 respectively).

**Fig 3.**
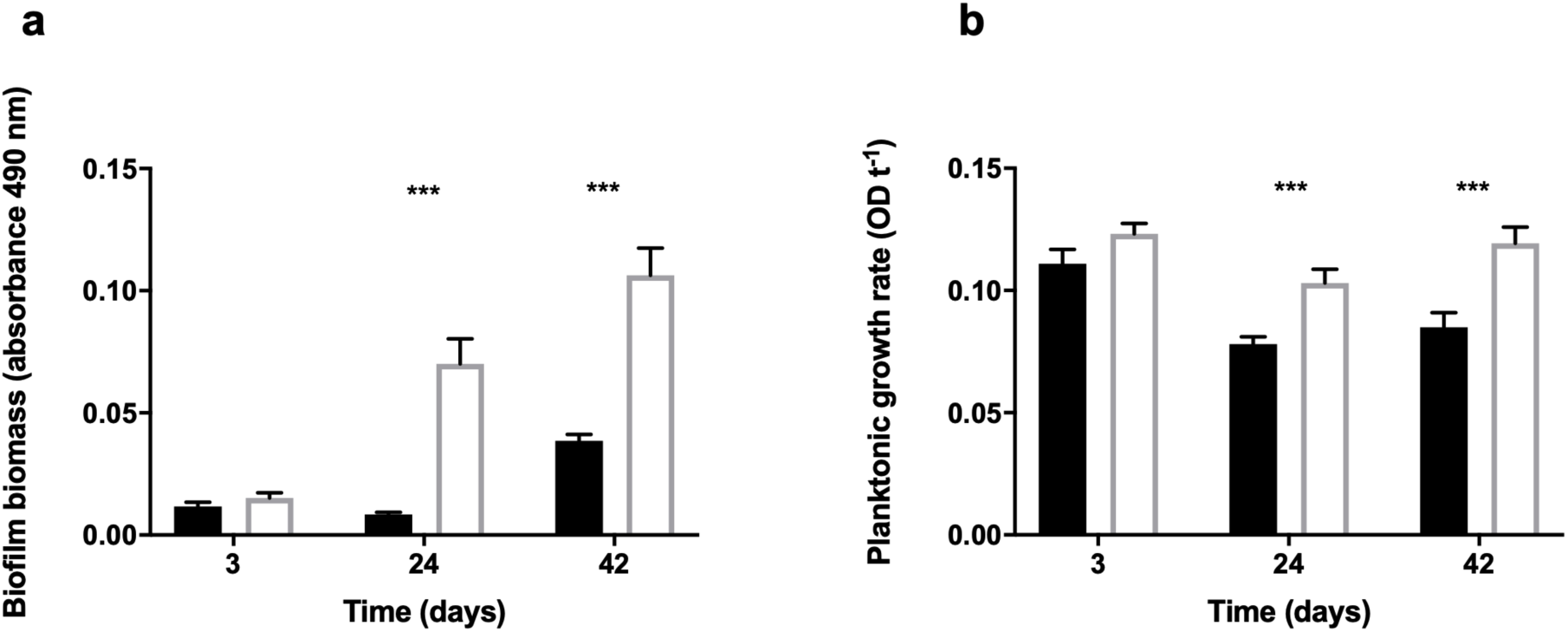
Effect of co-incubation of *P. aeruginosa* with *A. castellanii* on biofilm formation and growth. Biofilm biomass (a) and planktonic growth rates (b) of *P. aeruginosa* amoeba-adapted (closed) and non-adapted (open) isolates obtained from 3, 24 and 42 d populations when grown in LB_10_ media at 37°C. Data are presented as Means ± SEM. *** p < 0.001.

### Quantification of pyoverdine and rhamnolipids

*P. aeruginosa* isolates obtained after co-incubation with *A. castellanii* produced reduced quantities of pyoverdine compared to isolates from non-adapted populations (Fig 4a; *F*_1, 174_ = 45.74, *p* < 0.001). Although pyoverdine production was reduced in both amoeba-adapted and non-adapted populations (*F*_2, 174_ = 12.08, *p* < 0.001), the concentration of pyoverdine in supernatants from amoeba-adapted isolates from 3 d populations was reduced 2-fold compared to non-adapted isolates (*p* < 0.001) and was further reduced after 24, and 42 days of selection (*p* < 0.001).

**Fig 4.**
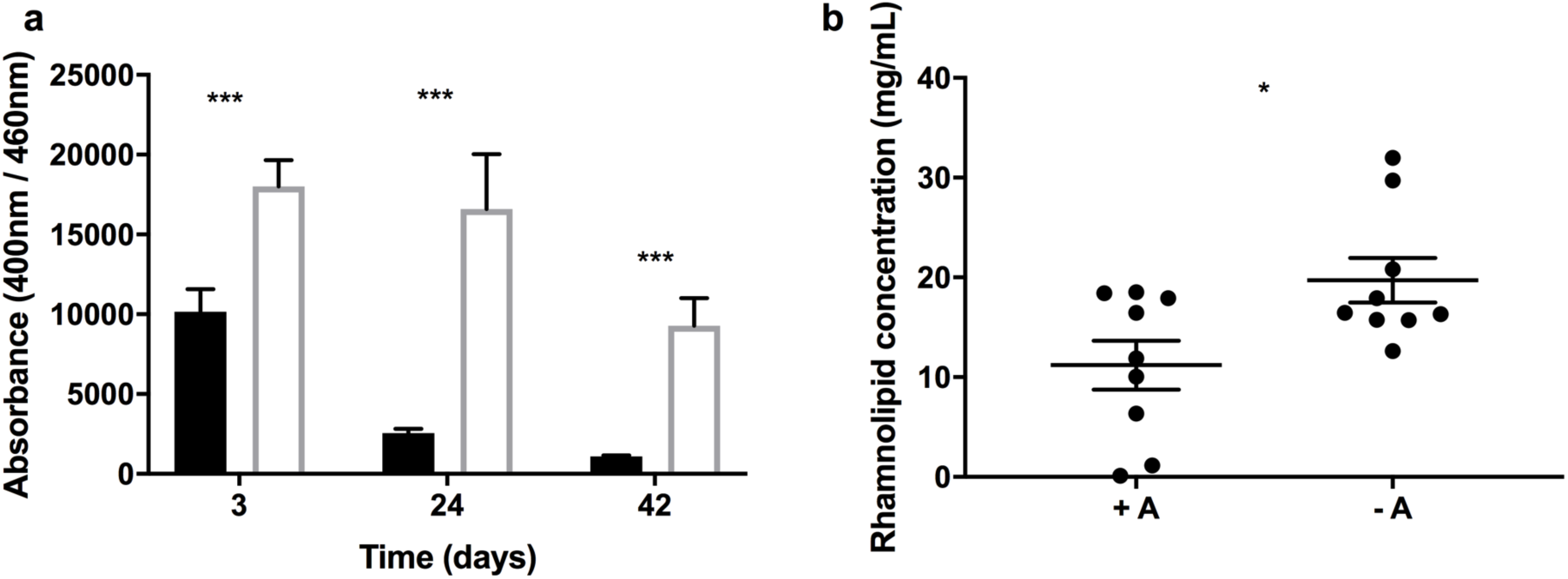
Effect of co-incubation of *P. aeruginosa* with *A. castellanii* on pyoverdine and rhamnolipid production. Quantification of pyoverdine (a) in supernatants from *P. aeruginosa* isolates obtained from amoeba-adapted (closed) and non-adapted (open) populations from days 3, 24 and 42. Quantification of rhamnolipids (b) in supernatants from *P. aeruginosa* isolates obtained from day 42 amoeba-adapted (+A) and non-adapted (– A) isolates using the orcinol method, with a correction factor of 2.5. Data are presented as Means ± SEM. * p < 0.05 *** p < 0.001.

Rhamnolipid production varied within the 42 d amoeba-adapted and non-adapted isolates but amoeba-adapted isolates produced less rhamnolipid overall when compared with the non-adapted population (Fig 4b; t_16_ = 2.571, p = 0.0205).

### Amoeba-adapted *P. aeruginosa* are more competitive than non-adapted isolates when grown with amoeba

To investigate whether adaptation with amoeba confers a fitness advantage to *P. aeruginosa* when grown with amoeba, we mixed fluorescent-tagged amoeba-adapted and non-adapted isolates and grew them together with amoeba. After 48 h co-incubation with amoeba, the proportion of amoeba-adapted cells is always higher when both amoeba-adapted::GFP (Fig 5a; F_1,4_ = 95.27, p = 0.000617) and amoeba-adapted::mCherry (Fig 5b; F_1,4_ = 11.85, p = 0.0262) are competed with the reciprocally tagged non-adapted strain, compared to no amoeba controls.

**Fig 5.**
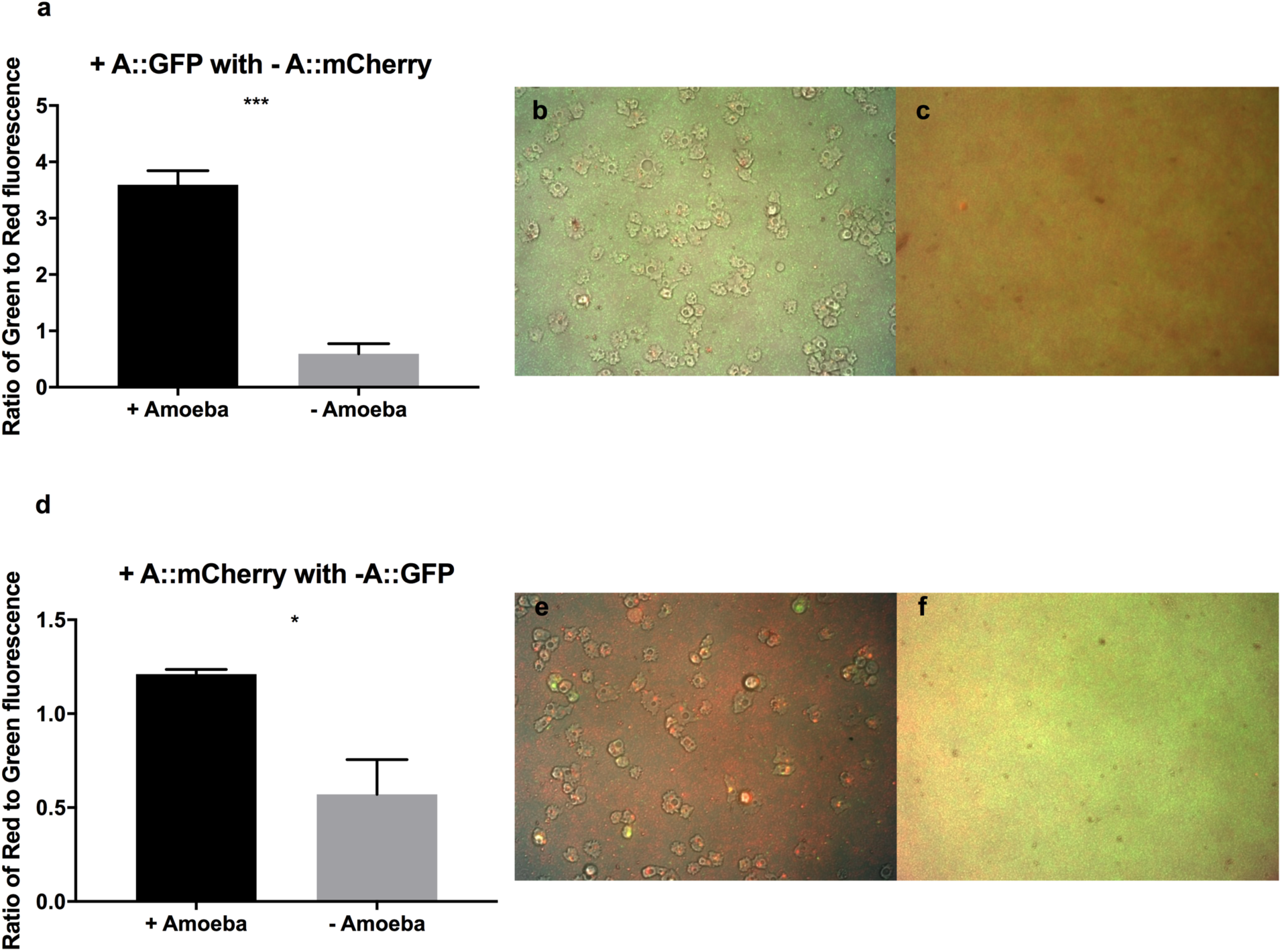
Competition of amoeba-adapted and non-adapted *P. aeruginosa* grown with amoeba. The fluorescence ratios of Day 42 + A::GFP *P. aeruginosa* mixed with – A::mCherry (a,b,c) and + A::mCherry with – A::GFP (d,e,f) after 48h of incubation with (black bars, b, e) and without (grey bars, c, f) *A. castellanii*. Data are presented as Means ± SEM. * p < 0.05, *** p < 0.001.

### Growth of amoeba-adapted and non-adapted *P. aeruginosa* in media or media supplemented with amoeba supernatant

To investigate whether the amoeba-adapted strains were utilizing amoeba secretions to out-compete the non-adapted strains, 9 randomly selected amoeba-adapted and non-adapted isolates were grown with and without the addition of amoeba supernatant to the growth media. The addition of amoeba supernatant to the growth media resulted in specific growth rate increases of −0.02 to 0.04 by 42 d amoeba-adapted and non-adapted *P. aeruginosa* isolates compared to the minimal media supplemented with the same amount of glucose. There was no significant difference in growth between amoeba-adapted and non-adapted isolates when grown in amoeba supernatant (Fig 6; t_12_ = 0.5147, p = 0.5147).

**Fig 6.**
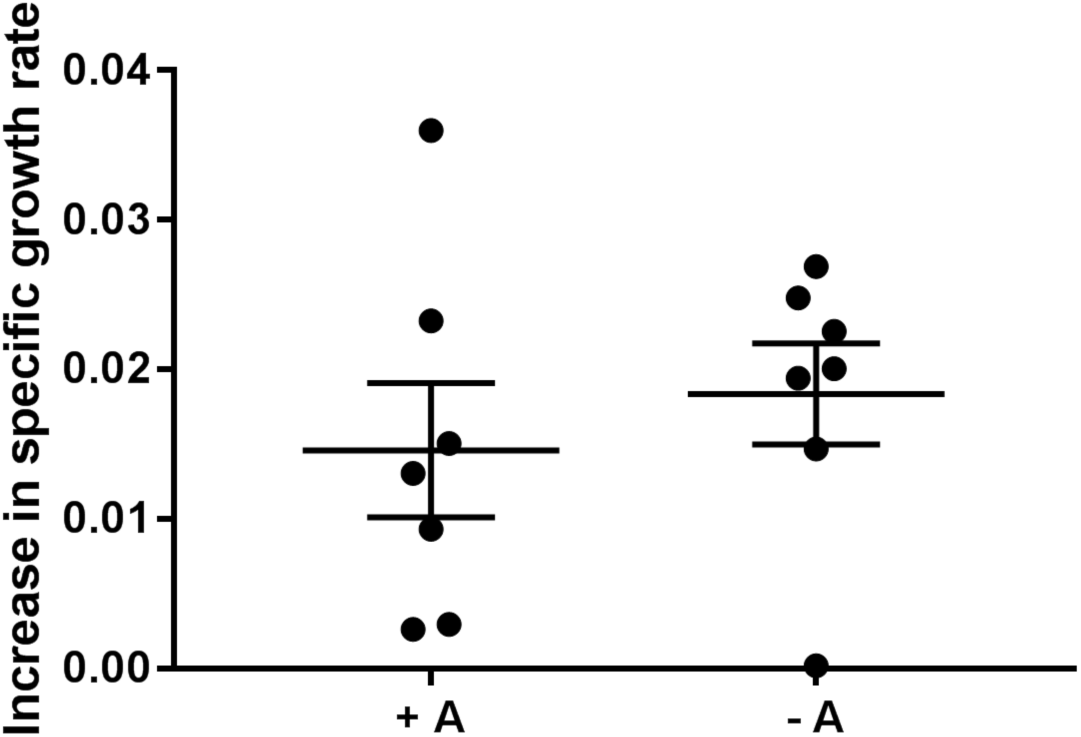
Growth of amoeba-adapted and non-adapted isolates in media supplemented with amoeba supernatant. Increase in specific growth rate of day 42 amoeba-adapted and non-adapted *P. aeruginosa* populations in M9 salts + 0.04% glucose supplemented with amoeba supernatant compared to no addition (n=9). Data are presented as Means ± SEM. No significant difference was observed in amoeba-adapted isolates.

### Amoeba-adapted *P. aeruginosa* isolates exhibit reduced uptake by and enhanced survival within amoeba and macrophages

Since the enhanced fitness of amoeba-adapted isolates in the presence of amoeba was not due to increased growth rate, the intracellular survival of 42 d amoeba-adapted and non-adapted populations were determined using a modified gentamicin protection assay. Intracellular CFUs 3 h after infection of non-adapted isolates within amoeba were higher than amoeba-adapted CFUs, however, after 24 h the numbers of surviving intracellular non-adapted cells had decreased and were comparable to the amoeba-adapted numbers (Fig 7a; Adaptation×Time F_1,32_ = 14, p < 0.001). The same trend was observed when the assay was conducted with raw 264.7 macrophages. There was a significant interaction of amoeba adaptation and incubation time (Fig 7b; Adaptation×Time F_4, 64_ = 6.692, p < 0.001), with a higher initial uptake of 42 d non-adapted populations compared to the amoeba adapted strains, resulting in higher initial intracellular CFU counts, followed by a constant decrease in viable intracellular numbers between 5 and 18 h post-infection. The 42 d amoeba-adapted populations were taken up by macrophage in lower initial numbers, and the number of viable intracellular CFUs did not decrease to the same extent as the non-adapted populations, resulting in comparable numbers at 18 h post-infection. At 24 h post-infection, macrophage cells infected with non-adapted *P. aeruginosa* exhibited morphological changes and appeared similar to those infected with the wild type strain (Fig 7c). Propidium iodide staining showed that many of these macrophages were dead. In contrast, macrophage infected with amoeba-adapted *P. aeruginosa* exhibited a more normal morphology, with fewer cells taking up the propidium iodide stain.

**Fig 7.**
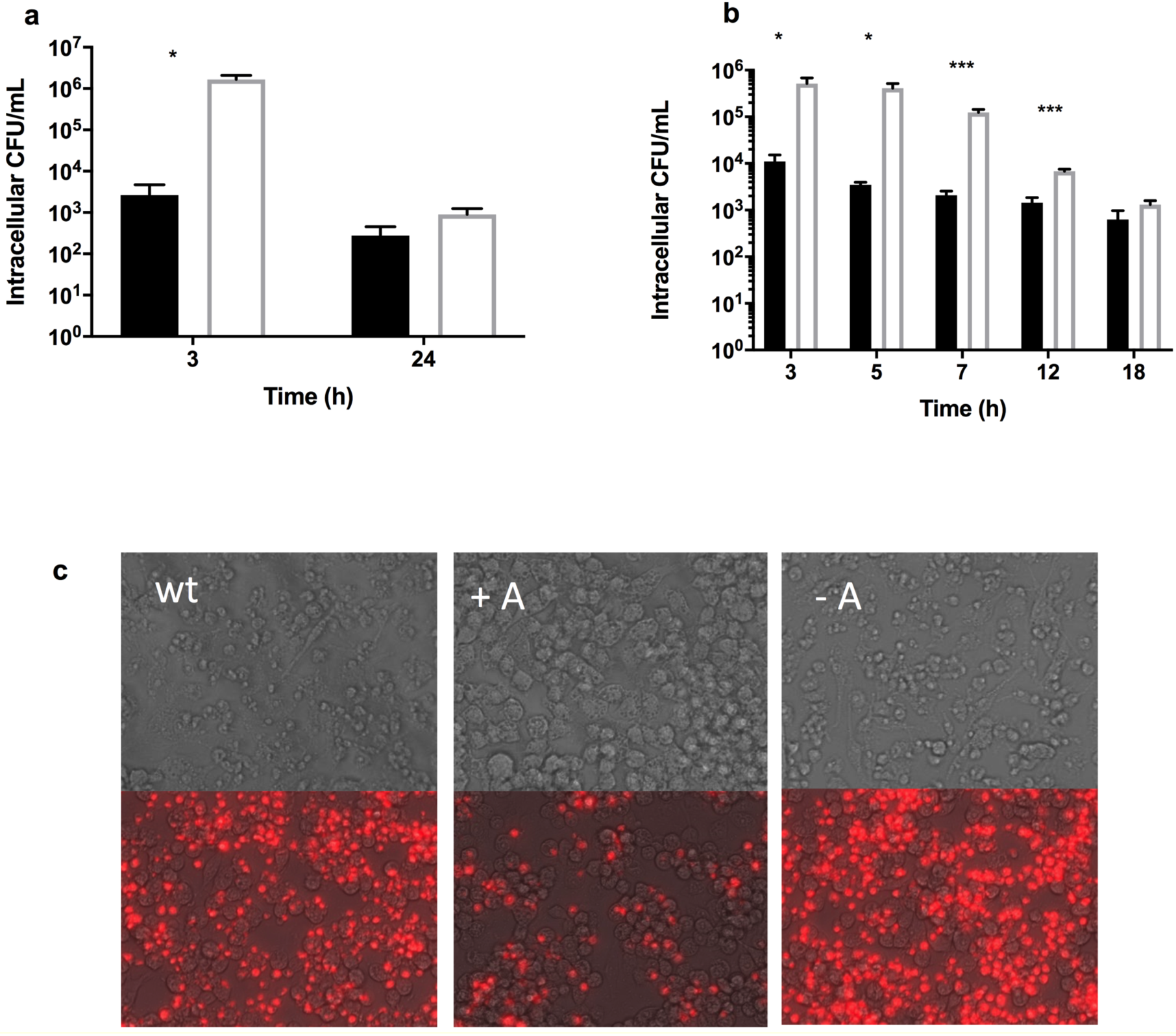
Intracellular survival of day 42 amoeba-adapted (closed) and non-adapted (open) strains in CFU ml^-1^ over time in a modified gentamicin protection assay (log-scale, n=3) conducted with (a) amoeba and (b) macrophage cells. Data are presented as Means ± SEM. *p < 0.05; *** p < 0.001. (c) Propidium iodide staining of raw 264.7 macrophage cells 24 h after infection with wild type, day 42 amoeba-adapted (A+) and non-adapted (-A) *P. aeruginosa*. Images are shown with and without the fluorescence to better illustrate changes in cell morphology.

### Amoeba-adapted *P. aeruginosa* isolates exhibit reduced uptake and enhanced survival in the presence of neutrophils

In order to determine if the intracellular survival of amoeba –adapted cells extends to other phagocytic cell types, we compared the ability of adapted and non-adapted *P. aeruginosa* to survive in the presence of human neutrophils. The internalization of 42 d amoeba-adapted and non-adapted populations were determined using a modified gentamicin protection assay. Intracellular CFUs 2 h after infection of non-adapted isolates within neutrophils were higher than amoeba-adapted CFUs (Fig 8a). Amoeba-adapted strains also had increased survival against human neutrophils when compared to non-adapted isolates (Fig 8b).

**Fig 8.**
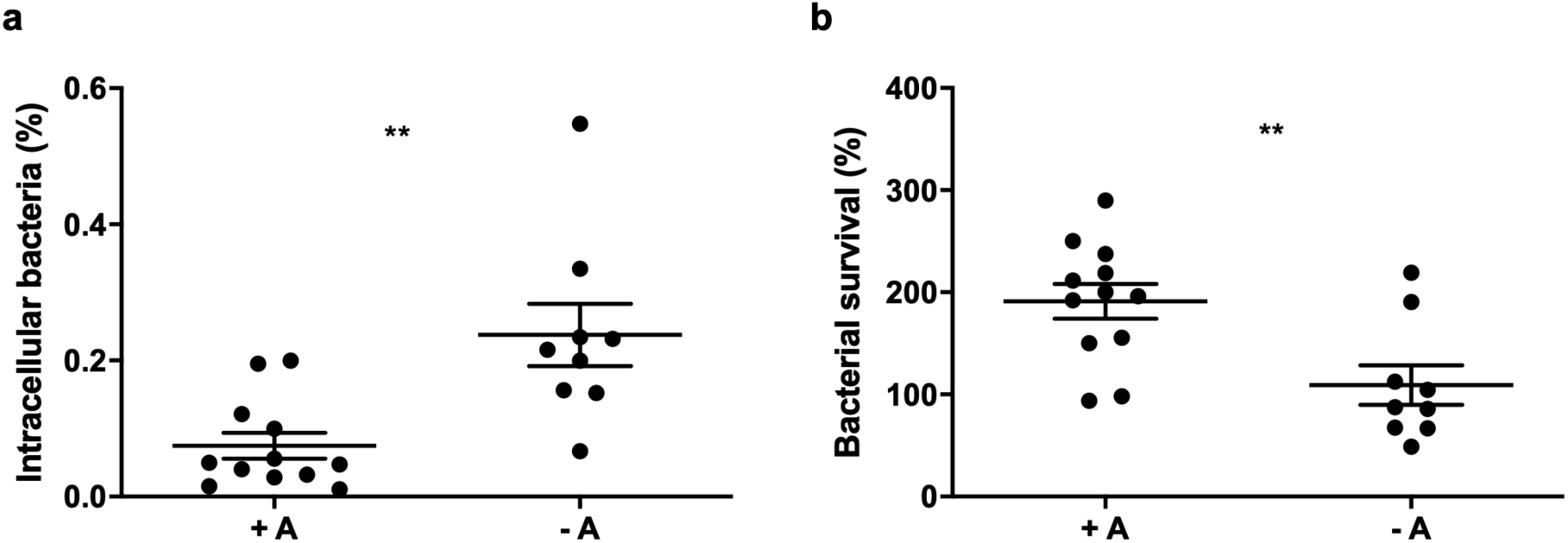
Amoeba-adapted *P. aeruginosa* exhibits reduced uptake by and enhanced survival in the presence of human neutrophils. Survival of day 42 day (+ A) amoeba-adapted (4) and (-A) non-adapted (3) strains following incubation with human neutrophils where bacterial (a) uptake and (b) survival were determined. Counts were performed in triplicate and results are the pooled means ± SEM from individual experiments using 3 different donors. Groups were analysed via student T test. **p<0.01

### *P. aeruginosa* populations co-incubated with *A. castellanii* show reduced virulence

Most of the phenotypes explored above play a role in the pathogenesis of *P. aeruginosa.* Therefore, we tested for virulence in *Caenorhabditis elegans* fast and slow kill assays. The isolates derived from amoeba-adapted populations after 3 d co-incubation were significantly more toxic to nematodes compared to non-adapted populations, although nematodes exposed to isolates from both populations had a median survival of 8 h (Fig 9a; (χ^2^ (1, n = 54) = 31.77, p < 0.001). *C. elegans* feeding on isolates from the 42 d non-adapted populations survived longer than when feeding on isolates from the day 3 populations. Furthermore, isolates from the 42 d amoeba-adapted populations were significantly less toxic to *C. elegans* compared to their non-adapted counterparts in both fast kill (Fig 9b; χ^2^ (1, n = 54) = 140.7, p < 0.0001) and slow kill assays (Fig 9c; χ^2^ (1, n = 54) = 23.52, p < 0.0001).

**Fig. 9.**
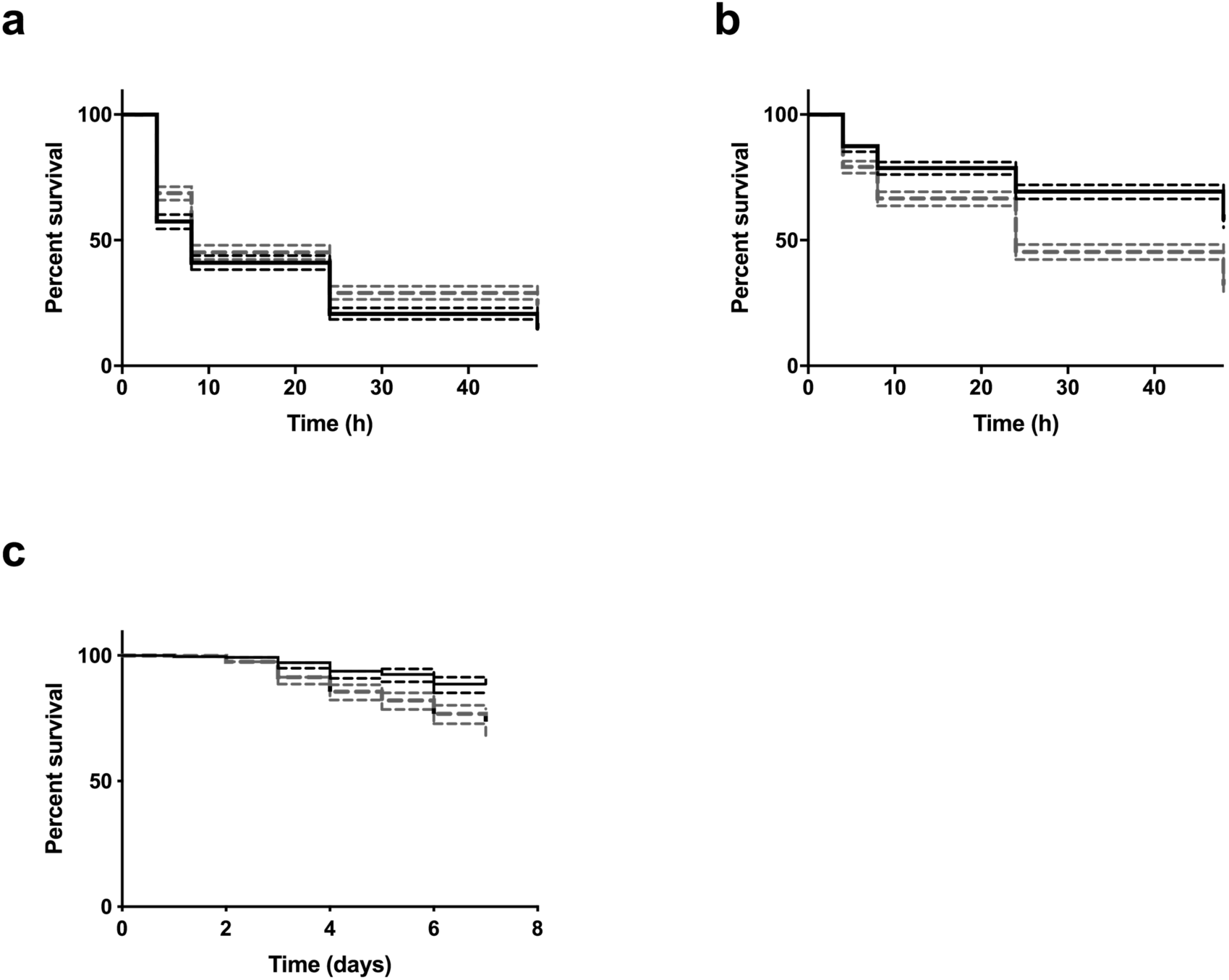
Effect of co-incubation of *P. aeruginosa* with *A. castellanii* in the *C. elegans* virulence assay. *C. elegans* survival curves after exposure to *P. aeruginosa* isolates derived from amoeba-adapted (black solid bars) or non-adapted (grey dashed bars) populations taken after 3 (a) or 42 (b) d for 0, 4, 8, 24 and 48 h in a fast kill assay. (c) Percent survival of *C. elegans* exposed to day 42 isolates of amoeba adapted and non-adapted *P. aeruginosa* populations over 7 days in a slow kill assay. Dotted lines indicate 95% CI.

## Discussion

Most opportunistic pathogens are not transmitted person to person but rather transit through the environment between hosts and therefore, it is likely that the environment plays a significant role in evolution of protective traits. Predation by protists is one of the major mortality factors for bacteria in the environment and it is likely that traits that protect against predation may also impact human hosts during infection (1–3). *P. aeruginosa* is responsible for a variety of nosocomial acute as well as chronic infections (15) in particular chronic lung infections in CF patients (16). Here we investigated the phenotypic and genotypic changes that occur during co-incubation of *P. aeruginosa* with *A. castellanii* for 42 days with a focus on virulence traits. Populations were collected on days 3, 24 and 42 and individual isolates were randomly collected at each time point.

### Motility, biofilm formation and secretion of secondary metabolites

Sequencing of populations obtained from the day 42 co-incubation revealed 54 and 65 nsSNPs in adapted and non-adapted populations, respectively. Gene functions of nsSNPs in coenzyme metabolism, lipid metabolism, cell motility and secondary metabolite production occurred solely within the amoeba-adapted population, while COGs representing inorganic ion transport and energy production were over-represented in the non-adapted population (Fig 1). Phenotypic assays of random isolates from the adapted and non-adapted populations confirmed loss of function in many traits associated with these genes. For example, amoeba-adapted isolates showed reductions in twitching and swimming motility from days 3 to 42 and a decrease in swarming motility for day 3 and 24 isolates, but not for the day 42 isolates (Fig 2). In addition, there was a decrease in planktonic growth rate and biofilm formation by amoeba-adapted isolates compared to non-adapted isolates (Fig 3). It is possible that the loss of biofilm is a result of selection against flagella or adhesive pili as these surface structures are immunogenic and act as ligands for phagocytes, including amoeba (21, 22). In fact, a large number of *P. aeruginosa* isolates from CF patients also lack flagella and pili and these mutants are resistant to phagocytosis by macrophage (23).

There were also a number of nsSNPs in genes relating to secretion of secondary metabolites in the adapted population. Phenotypic assays revealed that pyoverdine secretion was significantly reduced in the adapted isolates as early as day 3 and remained low throughout the later time points (Fig 4). Loss of pyoverdine in *P. aeruginosa* in chronic cystic fibrosis infections has also been reported (24). In addition, we observed an overall decrease in production of rhamnolipids, however, this was only a subset of the isolates tested. Unlike other traits that were largely or completely lost, decreases in rhamnolipid secretion were variable, implying that there may be a weaker selective pressure against this phenotype during co-adaptation.

### Uptake by and survival in amoeba and immune phagocytes

Results presented here reveal that amoeba-adapted isolates are taken up less readily and have increased intracellular survival in *A. castellanii* and RAW264.7 macrophage cells compared to non-adapted isolates, although the exact mechanisms are not known (Fig 5 and 7). Similarly, we saw reduced internalization of amoeba-adapted isolates by human neutrophils (Fig 8). The loss of immunogenic flagella and pili may contribute to reduced uptake (25), as well as a reduction in chemotaxis due to mutations in chemotaxis genes in all amoeba-adapted populations.

Protozoa have been reported to release dissolved free amino acids and other nutrients when grazing on bacterial prey (26). A reduction in chemotaxis would decrease the encounter of predator and prey since normally *P. aeruginosa* is attracted to amoeba where it attaches to the surface of the amoeba. Likewise changes in LPS also contribute to host avoidance, enhancing fitness of amoeba-adapted strains in the presence of amoeba or macrophage. Interestingly, PA3349 encoding a protein necessary for flagella-mediated chemotaxis was mutated in all amoeba-adapted lineages and is needed for acute but not for chronic *P. aeruginosa* infections (27, 28).

The finding that fitness gains from adapting to amoeba could be similarly conferred to interactions with macrophage cells, perhaps demonstrates the overlaps in traits used by *P. aeruginosa* to interact with both host types. Much work has been done to show that specialist intracellular pathogens such as *Legionella pneumophila* can evade the defenses of both amoeba and macrophage cells during endocytosis in order to form an intracellular replication niche (29). *P. aeruginosa* is more of an environmental generalist than an intracellular pathogen and has not been observed to form intracellular replication vacuoles, even though it has been recovered from within environmental amoeba (30). In addition, intracellular replication has been described in non-phagocytic epithelial cells and is dependent on a functional type-3 secretion system (31). In the amoeba co-adapted isolates, there was a general decrease rather than increase in intracellular cell numbers, which indicates that it is unlikely that *P. aeruginosa* is replicating intracellularly under the conditions used here.

This work also highlights the complexities of host-pathogen interactions and potential overlaps in these processes, even between cells as far diverged as mammalian macrophage cells and single-celled amoeba. The *A. castellanii* genome contains homologues of interferon-γ-inducible lysosomal thiol reductase enzyme (GILT), interferon inducible GTPase, and the NRAMP homologue, all of which play a role in antimicrobial defense in mammalian cells (32). More research is needed to determine whether these defenses play a role in *P. aeruginosa* infection and whether *P*. *aeruginosa* possesses mechanisms to evade such defenses in order to survive intracellularly within both phagocytic cell types.

### *P. aeruginosa* populations co-incubated with *A. castellanii* show reduced virulence

In order to investigate how constant predation pressure affects *P. aeruginosa* phenotypic and genotypic traits, we adapted *P. aeruginosa* to amoeba for 42 days. Interestingly, the amoeba-adapted isolates were less virulent in both fast and slow-kill *C. elegans* assays (Fig 9). Although the initial 3-day virulence levels were comparable, co-adaptation with amoeba for 42 days resulted in a loss of virulence against *C. elegans* by both fast kill and slow kill mechanisms. Fast and slow killing involves different virulence factors. For example, fast killing is usually due to the production and secretion of diffusible secondary metabolites, while slow killing occurs after colonization and infection of the gut (33). The production of hydrogen cyanide is the main virulence factor involved in the fast killing of *C. elegans* (34), while the regulated export of proteins is needed for slow killing (33).

Consistent with the reduction in *C. elegans* killing, we observed reductions in key virulence traits, such as motility, biofilm formation, and secondary metabolite production in days 24 and 42 amoeba-adapted isolates. The loss of motility and secondary metabolites probably contributes directly to reduced *C. elegans* mortality. In *P. aeruginosa*, type-IV pili are involved in adherence, swarming and twitching motility, and virulence. *P. aeruginosa* mutants deficient in *pilA* and *pilT* demonstrate reduced pathogenesis in mice compared to the wild type (35). Virulence towards nematodes has been partly attributed to pyoverdine. For example, when *Pseudomonas syringae* interacted with *C. elegans*, the genes, *pvdJ* and *pvdE* that are involved in the synthesis of pyoverdine, were significantly upregulated on fast kill agar (36). A similar response was observed in *P. aeruginosa*, where ‘red death’ type killing of *C. elegans* was shown to be partly due to the production of pyoverdine (37). Additionally, *P. syringae* Δ*pvdJ* and Δ*pvdL* mutants were unable to produce pyoverdine and an unrelated toxin, tabtoxin, demonstrated reduced AHL production and attenuated virulence against the tobacco plant host (38).

Rhamnolipids are quorum-sensing regulated secondary metabolites expressed in biofilms when neutrophils are present (39) that lyse macrophages, neutrophils and protozoans. If the gene is not expressed under the conditions of our co-adaptation, it may not be under strong negative selective pressures. Regulation of expression may be one mechanism that allows for the retention of certain virulence traits under predation.

### Commensalism as an adaptive strategy

The co-adapted isolates had mutations in many virulence phenotypes, including loss of motility, slower growth and increased intracellular survival. These adaptations represent a commensal strategy that may indicate adaptation towards co-existence with amoeba. *P. aeruginosa* consistently adopts a more commensal lifestyle during adaptation to different hosts. For example, experimental evolution experiments with *C. elegans* demonstrated that *P. aeruginosa* evolved an attenuated virulence phenotype after serial passages (12). This is in contrast to other pathogens where virulence has been demonstrated to increase after serial passages (40). The decrease in acute virulence phenotypes also occurs in *P. aeruginosa* strains isolated from chronically infected CF patients (17, 41, 42).

In co-evolution with *C. elegans*, *lasR* and *rhlR* quorum sensing mutations occurred early in the adaptation process. In addition, *lasR* mutations are prevalent across multiple CF isolates (43). The regulatory genes *lasR* and *rhlR* control the expression of many virulence genes, and *las* and *rhl* mutants have been shown to be less virulent in models of wound infection (44). It has been proposed that pleiotropic adaptive mutations in global regulatory genes are more likely to occur than multiple mutations in individual virulence traits. Mutations in *lasR* occurred in one of the three amoeba-adapted replicates, so stronger selection may be driving the loss of individual virulence traits in the other amoeba-adapted populations. In CF lineages, *P. aeruginosa* adaptations include gains in mucoidy and antibiotic resistance, and loss of secondary metabolites and motility (17), of which the latter two were observed in the amoeba-adapted populations. The parallel losses of secondary metabolites and motility are striking and evoke the question of whether the selective forces driving these traits are the same in both systems.

Advantageous traits may be lost if the cost of maintaining the traits outweigh the benefits. Loss of pyoverdine in *P. aeruginosa* CF infections has been shown to be driven by social selection (45). Pyoverdine is a strong scavenger for iron and non-producing cheats retain the pyoverdine receptor and are able to uptake iron. Only when extrinsic pyoverdine is completely lost do mutations appear in the receptor genes. In our study, although there were many *pvd* mutations, we did not observe any mutations in the receptor genes, suggesting that social selection may also be at play here. It is also possible that other avenues of iron uptake are preferentially utilized (46), as iron uptake occurs via *hemO* in late CF strains (47).

Strong negative selection against traits could also occur due to host recognition and the need for evasion by pathogens. Flagellin is the site of recognition by mammalian toll-like receptors, resulting in immune activation (22), and the site where *A. castellanii* binds *P. aeruginosa* before it is endocytosed (21). Additionally, flagellin and TLR-independent loss of motility has been shown to significantly reduce phagocytic uptake by mammalian immune cells (48), and may explain why loss of motility is such a strong selection factor in the co-adaption study presented here and in CF lineages. Several findings presented here support predator avoidance as a selection pressure. The parental *P. aeruginosa* strain exhibits chemotaxis and rapidly swims towards and are taken up by amoeba and macrophage cells within seconds of exposure. However, amoeba-adapted isolates do not attach to the surface of the amoeba. This is supported by experiments showing reduced uptake of amoeba-adapted populations by amoeba and macrophage. Loss of flagella and motility is therefore adaptive for the purpose of predator avoidance. Additionally, many chemotaxis mutants were detected in the population genomic data. For example, *pctB* and PA3349 were mutated in all three independent experimental amoeba-adapted populations. *pctB* is a methyl-accepting chemotaxis protein with a high affinity to glutamine (49), which has been shown to be involved in chemotaxis to wounded airway epithelial cells (50). The loss of chemotaxis in the amoeba-adapted strains would be consistent with predator avoidance.

In this study, it has been demonstrated for the first time that adaptation to a more commensal lifestyle may also confer benefits in an infectious context for a generalist pathogen, for it is clear that although amoeba-adapted cells are less virulent, they are still capable of invading and colonizing the *C. elegans* model. Similarly, adapted CF strains have been shown to be equally as capable as environmental strains of infecting a new host in a mouse model (51). Although amoeba may be thought of as training grounds for the formation of virulence traits, they may also be grounds for a more ‘chronic’ co-existence.

## Materials and methods

### Organisms and growth conditions

*P. aeruginosa* strain DK1 used for this study was initially obtained from a Danish CF patient (P30M0) (18). Unless otherwise stated, *P. aeruginosa* DK1 strain and population-derived isolates were grown in 10 mL lysogeny broth (LB_10_, BD Biosciences, USA) at 37 °C with shaking at 200 rpm. *A. castellanii* was obtained from the American Type Culture Collection (ATCC 30234) and was routinely maintained axenically in peptone-yeast-glucose (PYG) medium (20 g protease peptone, 5 g yeast extract, and 50 mL 2 M glucose L^-1^) at room temperature. Prior to use in experiments, *A. castellanii* was passaged and washed twice with 1 × phosphate buffered saline (PBS; Sigma-Aldrich, USA) solution to remove PYG media. *C. elegans* N2 Bristol was maintained on nematode growth medium (NGM) (per liter; 2.5 g Bacto-Peptone (BD Biosciences, USA), 3 g NaCl, 7.5 g agar, 1 mL 5 mg mL^-1^ cholesterol, 1 mL 1 M MgSO_4_, 1 M CaCl_2_, and 25 mL 1 M potassium phosphate buffer at pH 6) fed with *E. coli* OP50.

RAW 264.7 macrophage cell lines (ATCC TIB-71) were grown in Dulbecco’s modified Eagle Medium (DMEM; Thermo Fisher, USA) with 10 % fetal bovine serum (FBS) at 37 °C with 5 % CO_2_. Before use, cells were washed with PBS and treated with trypsin briefly before gentle detachment by scraping. Cells were then centrifuged at 1000 × *g* for 1 min and resuspended in experimental media before use.

### P. aeruginosa and A. castellanii co-incubation

Overnight cultures of *P. aeruginosa* grown in was centrifuged at 4000 × *g* for 5 min and washed twice with 1 × M9 salts solution (Sigma-Aldrich, USA; per litre, 6.78 g Na_2_HPO_4,_ 3 g H_2_PO_4,_ 1 g NH_4_Cl, 0.5 g NaCl). *A. castellanii*, at a concentration of 1 × 10^3^ cells mL^-1^, was seeded onto the surface of 25 cm^2^ tissue culture flasks with 0.2 µm vented caps filled with 10 mL 1 × complete M9 salts + 0.01% glucose. To maintain a strong selective pressure from amoeba, 100 µL of *A. castellanii* and *P. aeruginosa* was taken from percussed, 3-day-old established flasks and added to new flasks containing *A. castellanii* every 3 d. Three independent populations of *P. aeruginosa* with *A. castellanii* were established (amoeba-adapted populations).

In parallel with the co-incubation experiment, *P. aeruginosa* were maintained without *A. castellanii*. Briefly, *P. aeruginosa* was diluted to a cell concentration of 1 ×10^2^ cells mL^-1^ and added to tissue culture flasks containing 10 mL 1 × complete M9 + 0.01 % glucose. From these flasks, 100 µL of 3-day-old established *P. aeruginosa* culture was added to flasks containing fresh media every 3 d. Three independent populations of *P. aeruginosa* were established (– A populations).

Prior to re-inoculation and sampling, the biomass of each population was measured by spectrophotometry (Infinite^®^ M200, Tecan, Switzerland) at 600 nm. The number of *A. castellanii* within each amoeba-adapted population were enumerated using inverted microscopy (Zeiss Axio Observer.Z1 inverted wide-field).

### Isolation of intracellular *P. aeruginosa* populations

On days 3, 24 and 42, flasks containing *A. castellanii* (amoeba-adapted) were percussed until the amoebae detached and 1mL of the culture media was filtered through a 3 µm cellulose acetate membrane (Merck, Germany) to retain the *A. castellanii. A. castellanii* were resuspended in 5 mL of 1 × M9 salts and pelleted at 4000 × g for 5 min before resuspension in 100 µL 1 × M9 salts solution. *A. castellanii* were lysed by the addition of 100 µL of 1 % Triton –X for 1 min, the mixture was then pelleted and washed twice with 900 µL of 1 × M9 salts. The cell pellet was resuspended in 1 mL of 70 % LB_10_ + 30 % glycerol and stored at −80°C. The same treatment was applied to the *P. aeruginosa* cells from the non-adapted populations.

### Sequencing of *P. aeruginosa* populations and computational tools

*P. aeruginosa* populations were sequenced to determine genotypic changes that occurred in response to co-adaptation with amoeba. Genomic DNA was extracted from the parental wild type strain, and each amoeba-adapted and non-adapted population derived from days 3 and 42 using the QIAamp DNA mini kit (Qiagen, Venlo, Netherlands) according to manufacturer’s instructions. Sequencing libraries were prepared using the TruSeq DNA sample preparation kit (Illumina, San Diego, CA, USA), and sequenced on a MiSeq (Illumina, USA).

Reads were aligned to the *P. aeruginosa* PAO1 reference genome using CLC Genomics Workbench 9 (CLC Bio, Aarhus, Denmark). SNPs and small insertions and deletions (indels) were called using the probabilistic variant detection analysis, and mutations that were also in the parental strain were filtered out. Genes that contained SNPs in > 25% of the total gene reads were selected for functional analysis using the Database for Annotation, Visualization and Integrated Discovery (DAVID) v6.8 functional tools (52, 53). Gene lists were uploaded onto the DAVID website (https://david.ncifcrf.gov/) and annotations were limited to *P. aeruginosa*. A functional annotation table was compiled for amoeba adapted populations obtained from day 42 of the experiment (Table 1).

### Isolation of adapted strains

To facilitate further phenotypic analysis, *P. aeruginosa* cells obtained from amoeba-adapted and non-adapted populations on days 3, 24 and 42 were plated onto LB_10_ agar and incubated overnight at 37°C. Ten colonies were randomly selected from each population using a numbered grid and a random number generator. These isolates were stored in 1 mL 70 % LB_10_ + 30 % glycerol at −80°C.

### Assessment of surface colonization and planktonic growth

To determine if adaptation with *A. castellanii* altered surface colonization by *P. aeruginosa*, the biomass of attached cells was quantified by crystal violet staining as previously described (54). Briefly, overnight cultures of each isolate were added to 96 well plates containing 100 µL 1 × M9 + 0.4 % glucose. Plates were incubated at room temperature with agitation at 80 rpm for 24 h. To separate the suspended biomass from the attached biomass, each well was washed once with 1 × PBS. Cells attached to the plate surface were stained with 200 µL of 0.3 % crystal violet and incubated for 20 min, after which unbound crystal violet was removed by washing 3 times with 1 × PBS. Crystal violet was liberated from the cells with 300 µL of 10 mL of absolute ethanol. The absorbance of the crystal violet was measured with a spectrophotometer at 590 nm (Infinite^®^ M200, Tecan, Switzerland) in triplicate.

The planktonic growth rate of each isolate was assessed by adding log phase bacterial cultures to 96 well plates containing 200 µL of 1 × M9 + 0.01 % glucose and incubating with agitation at 80 rpm for 18 h at 37°C. The suspended biomass within each well was transferred to a fresh plate and measured by spectrophotometry (OD _600 nm_) (Infinite^®^ M200, Tecan, Switzerland). The specific growth rate was determined by applying the formula µ = 2.303 ((log OD2) – (log OD1)/(t2-t1)). Assessment of the planktonic growth rate was performed in triplicate for each isolate, n = 30 for each time and treatment.

### Motility assays

Twitching, swarming and swimming motility were assessed as previously described, using motility agar (20 mM NH_4_Cl, 12 mM Na_2_HPO_4_, 22 mM KH_2_PO_4_, 8.6 mM NaCl,1 mM MgSO_4_, 100 µM CaCl_2_, 2 gL^-1^Dextrose, 5 g L^-1^casamino acids) containing 1, 0.5 or 0.3 % wt vol^-1^ agarose (Bacto™, BD Biosciences, USA), respectively (55, 56). Five milliliters of motility agar were added into the wells of 6 well plates and dried under laminar flow for 1 h. Isolates were inoculated into the center of the well using 10 µL pipette tips, either to the base of the plate for assessment of twitching motility or mid-agar for assessment of swimming and swarming. Twitching and swarming plates were incubated at room temperature for 48 h and swimming plates were incubated for 24 h prior to imaging with a digital camera (Canon EOS 600D digital single-lens reflex (DSLR) mounted on a tripod, to allow for phenotypic characterization of the resulting colonies and comparative endpoint twitch, swarm and swim distances. Determination of the zone of motility was semi-quantitatively analyzed using ImageJ image analysis software. Motility was assessed in triplicate (n = 3).

### Quantification of pyoverdine

To determine if adaptation with amoeba (3, 24 and 42 d) affects the production of pyoverdine, isolates were grown overnight in 1 mL LB_10_ media. Cells were removed by centrifugation at 5200 × *g* for 5 min and the absorbance of the supernatant was determined with a spectrophotometer (Infinite^®^ M200, Tecan, Switzerland) at excitation 400 nm and emission 460 nm in triplicate.

### Quantification of rhamnolipids

The orcinol method (57) was used to quantify the production of rhamnolipid biosurfactant of nine randomly selected adapted and non-adapted isolates from the day 42 population. Briefly, overnight *P. aeruginosa* LB cultures were diluted to OD_600_ 0.01 in 25 mL of AB minimal media (58) supplemented with 2 g glucose and 2 g casamino acids L^-1^ and grown overnight at 37 °C with shaking at 200 rpm. The cell density was determined (OD_600 nm_) before filtration and extraction of crude rhamnolipid from the supernatant two times using diethyl ether (7 mL). The organic layer was collected, combined, and concentrated in a vacuum concentrator (SpeedVac, Thermo Scientific) at 0 °C for 1 h followed by 2 h at 25 °C, until white solids formed. The solids were resuspended in 500 µL of water and 50 µL of this solution was mixed with 450 µL of freshly prepared orcinol (0.19 % in 53 % H_2_SO_4_). Samples were incubated at 80 °C for 30 min and allowed to cool at room temperature for 15 min before quantification of absorbance (OD_421_). The absorbance was normalized to cell concentration (OD_600 nm_) for each sample and a factor of 2.5 was applied to convert values from a rhamnose standard curve to rhamnolipid concentration (59).

### Nematode survival assay

To determine if *P. aeruginosa* adaptation to *A. castellanii* altered bacterial virulence, we tested the survival of *C. elegans* sp. Bristol N2 after feeding on *P. aeruginosa*. Axenic *C. elegans* were obtained via the egg-bleach synchronization method, plated onto NGM agar and fed with heat-killed *Escherichia coli* OP50. L4 stage worms were re-suspended in 1× M9 salts solution and 10-30 worms were drop plated onto 35 mm dishes containing 2 mL fast or slow kill agar (33) containing lawns of pre-established *P. aeruginosa* obtained from amoeba-adapted or non-adapted populations. Plates were incubated at 22 °C and worm numbers were scored by microscopy at 0, 4, 8, 24 and 48 h for fast kill assays, and once per day for slow kill assays. Nematode toxicity was tested using 9 randomly selected *P. aeruginosa* isolates from each treatment and from times 3 and 42 d. Nematode survival assays were repeated twice independently, and each experiment was performed in triplicate.

### Fluorescent tagging of isolates

To prepare fluorescently-tagged strains of *P. aeruginosa* obtained from amoeba-adapted and non-adapted populations, two isolates were randomly selected from the 42 d population and grown overnight at 37 °C in LB broth. Electroporation was performed as previously described (60). One milliliter of *P. aeruginosa* at a cell density of 1 × 10^8^ cells mL^-1^ was pelleted and washed twice with 300 mM sucrose. The expression tag carrying plasmid pUC18-TR6K-mini-Tn7T-Gm-GFP (0.5 µg) (expresses a green fluorescent protein *gfp*; emission 488 nm/excitation 509 nm) (61) or pUC18T-miniTn7T-Gm-Mcherry (0.5 µg) (expresses a red fluorescent protein mCherry emission 587 nm/excitation 610 nm) (62) was mixed with 1 µg pTNS1 helper plasmid and 300 µl resuspended bacteria, in a 2-mm-gap electroporation cuvette. This mixture was electroporated at 1.8kV/25µF/2100 Ω, 2.5kV cm^-1^ in a Gene Pulser apparatus (BIO-RAD, Hercules, CA, USA). Cell recovery was performed in ice-cold super optimal broth with catabolite repression media (SOC) (10 mM NaCl, 2.5m M KCl, 10m M MgCl_2_, 10m M MgSO_4_, 20 g L^-1^ tryptone, 5 g L^-1^ yeast extract + 2% glucose. Cells were incubated with shaking for 3h at 37 °C. One hundred microliters of culture were plated onto LB_10_ plates supplemented with 200 ug mL^-1^ gentamicin to select for GFP and mCherry transformants. Bacterial stocks were derived from a single transformed colony.

### Competition assays

To determine if prior exposure of *P. aeruginosa* to amoeba increased competitiveness when grown with amoeba, a competition assay was performed. One isolate from the 42 d amoeba adapted population and one isolate from the 42 d – A population were fluorescently tagged as previously described. *A. castellanii*, at a cell concentration of 1 × 10^5^ cells mL^-1^ was added to 24 well plates (Falcon) in 450 µL 1 × M9 salts + 0.01 % glucose solution. Overnight cultures of GFP or mCherry-labelled *P. aeruginosa* were grown in LB_10_ broth supplemented with 200 µg gentamicin at 37 °C with agitation at 200 rpm. The amoeba adapted::gfp and – A::mCherry isolate or the amoeba adapted::mCherry and the – A::gfp isolate were mixed in equal proportion and added to the wells containing amoeba to a final bacterial cell concentration of 2 × 10^6^ cells mL^-1^. Each experiment was conducted in triplicate. The plates were incubated at room temperature with agitation at 60 rpm for 48 h before imaging on a Zeiss Z1 inverted wide field microscope. Acquired images were deconvoluted in Autoquant X3 (Bitplane) before quantification of the relative red and green fluorescence in the field of view using Imaris 8 (Bitplane).

### Growth of *P. aeruginosa* populations in amoeba supernatant

To investigate whether fitness differences were due to enhanced growth by utilizing resources released by amoeba, nine randomly selected day 42 adapted and non-adapted *P. aeruginosa* isolates were grown in media supplemented with 50 % amoeba supernatant in order to compare their growth rates. Amoeba supernatant was obtained by growing *A. castellanii* in M9 salts minimal media + 0.04 % glucose with heat killed *P. aeruginosa* (made by incubating overnight cultures at 65 °C for 3 h and plating on LB agar to check that no live cells remain) for 3 d, and then filtering the suspension through an 0.22 µm filter (Pall, USA). Overnight cultures of *P. aeruginosa* isolates grown in LB were adjusted to the same optical density (OD_600 nm_) and added to amoeba supernatant or M9 salts + 0.04 % glucose. Planktonic growth rates were quantified from optical density readings (OD_600 nm_) using a Tecan microplate reader as previously described.

### Uptake and intracellular survival of *P. aeruginosa* populations in macrophages

To investigate the dynamics of uptake and intracellular survival of 42 d +A and −A *P. aeruginosa* populations within macrophages, overnight LB cultures of adapted and non-adapted *P. aeruginosa* populations were added to RAW264.7 macrophage cells (5 × 10^4^ cells/well in 96-well tissue culture plates) in DMEM without FBS at a multiplicity of infection (MOI) of 100:1. The infected cells were incubated at 37 °C with 5 % CO_2_. After co-incubation for 1 h, the media was removed and replaced with media containing 100 µg mL^-1^ gentamicin to kill extracellular bacteria. Macrophage were washed with PBS and lysed at 3, 5, 7, 12, and 18h post-infection and CFU counts were performed to enumerate surviving intracellular cells. Propidium iodide (ThermoFisher LIVE/DEAD Cell Viability kit) staining was done to determine the state of the host cells 24h post-infection.

### Uptake and survival of *P. aeruginosa* populations in the presence of neutrophils

*P. aeruginosa* (4 adapted and 3 non-adapted) strains from overnight culture were washed once in PBS then diluted in PBS (OD=0.1, ∼1×10^8^) and resuspended in complete media (RPMI + 2% heat inactivated autologous plasma) to experimental concentrations just prior to infection. Neutrophils were isolated from whole blood, collected from healthy donors in lithium heparin vacutainer tubes and separated using polymorphprep (axis shield) and centrifugation. RBCs were hypotonically lysed and neutrophils washed in HBSS (without Ca^+^ or Mg^+^). Neutrophils were counted and resuspended at their final concentration in complete media. In a 96-well plate, neutrophils were added to wells for challenge (neutrophil+) and complete media added to control wells (neutrophil-). *P. aeruginosa* was added to both PMN+ and PMN-wells at a MOI of 100:1 and incubated for 1 h at 37°C, 5% CO2. After co-incubation for 1 h bacterial survival was determined by serial dilution and plating on LB for enumeration. Uptake was determined by media removal and replacement with media containing 100 µg mL^-1^ gentamicin to kill extracellular bacteria. At the experiment endpoint a sample of infection was taken and lysed in a new 96 well plate, followed by serial dilution and plating onto LB agar. CFUs were determined by counting and percent inoculum determined as [CFU of neutrophil + wells/CFU neutrophil-wells x 100]. Counts were performed in triplicate and results are the pooled Means ± SEM from individual experiments using 3 different donors.

### Human Ethics

Ethics for whole blood collection was obtained from the University of Wollongong Human research Ethics Committee (HREC # 08/250).

### Statistical analyses

Phenotypic differences between *P. aeruginosa* isolates obtained from amoeba-adapted and non-adapted populations at specific time points (3, 24 and 42 d) were determined by ANOVA, with amoeba adaptation (with or without *A. castellanii*) as a fixed factor and adaptation time (3, 24, 42 d) as a random factor. Multiple testing was conducted using the Tukey Post-hoc Test. All phenotypic data were log transformed (ln (x+1)) prior to analysis to improve normality. *P*-values < 0.05 were considered significant. Nematode survival curves were constructed with GraphPad Prism v 6.0 using the Kaplan-Meier method. Differences between nematode survival after exposure to *P. aeruginosa* isolates from amoeba-adapted or non-adapted populations were determined using log-rank tests with significance given to *p-*values < 0.05. Differences between neutrophil uptake and survival of amoeba-adapted and non-adapted strains were analyzed via student’s T tests.

## Acknowledgments

The authors would like to acknowledge support from The ithree Institute at the University of Technology Sydney, Sydney, Australia, the Australian Research Council Discovery Project (DP170100453) and by the National Research Foundation and Ministry of Education Singapore under its Research Centre of Excellence Program to the Singapore Centre for Environmental Life Sciences Engineering, Nanyang Technological University.

